# aaKomp: Alignment-free amino acid *k*-mer matching for genome completeness assessment at scale

**DOI:** 10.64898/2026.03.19.713078

**Authors:** Johnathan Wong, Lauren Coombe, René L Warren, Inanc Birol

## Abstract

In *de novo* sequencing projects, genome assembly optimization requires evaluating a number of candidate assemblies to identify optimal tool parameters. Yet, current completeness assessment tools like BUSCO and compleasm require 10-80 minutes per evaluation for gigabase-scale genomes, transforming what should be rapid iteration into time-intensive processes. These tools rely on alignment-based approaches and fixed ortholog databases, limiting their scalability across the tree of life. We present aaKomp, a scalable alignment-free tool that leverages amino acid *k*-mer matching and multi-index Bloom filters for rapid genome completeness assessment. Unlike current utilities, aaKomp supports user-defined reference databases, enabling customized assessments for any organism. In benchmarking against state-of-the-art tools using simulated T2T-CHM13 datasets, aaKomp achieved 68-fold faster execution and 15-fold lower memory consumption while maintaining accuracy. Testing on 94 Human Pangenome Reference Consortium assemblies and a European Eel assembly, aaKomp maintained one-minute runtimes (1.2 ± 0.35 min) and low memory usage (<13.64 GB). aaKomp’s scoring system provides nuanced estimates rather than threshold-based classifications, offering increased resolution for tracking incremental improvements during iterative workflows. aaKomp’s speed, memory efficiency, and flexible database generation makes it well-suited for modern and biodiverse projects requiring the evaluation of hundreds of assemblies.

## Introduction

Advances in genome sequencing, including improvements in throughput and read lengths, have driven a substantial increase in the number of genome assemblies generated across a wide range of species (Satam et al. 2023; Sayers et al. 2025; “zyndagj/ncbi_genomes” n.d.). However, producing a high-quality genome assembly remains a computationally complex task, requiring optimization of multiple interdependent objectives, such as contiguity, accuracy, and completeness, through careful algorithm and parameter selection, followed by evaluation of the resulting assemblies for contiguity, base accuracy, and functional completeness (Rhie et al. 2021; Li and Durbin 2024). No single assembly pipeline or configuration performs optimally across all datasets, which vary in data type and quality, and represent genomes of different sizes, complexities, and repeat content (Jackman et al. 2017; Kolmogorov et al. 2019; Shafin et al. 2020; Cheng et al. 2021; Wong et al. 2023a). As a result, researchers often generate multiple candidate assemblies per sample, followed by iterative tuning to refine assembly quality. This process, combined with the growing scale and diversity of genome assemblies, has created a need for fast, scalable methods to evaluate assembly completeness.

This challenge is exacerbated in large-scale initiatives like the Human Pangenome Reference Consortium (HPRC) (Liao et al. 2023) and the Earth BioGenome Project (Lewin et al. 2018), which aim to generate thousands of reference-grade genomes spanning global genetic diversity. In these contexts, efficient and reliable evaluation of assembly quality becomes essential for selecting, and refining genome builds. Common assembly evaluation metrics include contiguity statistics, such as NGA50, which measures the length at which half of the genome is covered by alignment blocks of that size or longer, and base-level accuracy estimates, such as assembly consensus quality value (QV), which represents a log-scaled probability of error for the consensus base calls (Mikheenko et al. 2018; Rhie et al. 2020). These approaches are not always feasible for non-model organisms without an established reference or matched high-quality *k*-mer data, usually from short reads, and are biased by alignments to the reference genome for model organisms.

In addition to contiguity statistics and assembly consensus QV, assessing functional completeness—i.e., whether an assembly contains the expected set of genes—offers a biologically meaningful measure of quality, with higher completeness indicating a more comprehensive representation of the coding genome. This is particularly important when reference genomes are unavailable, making gene content-based methods a preferred practice, especially for non-model organism genome assembly workflows. BUSCO (Benchmarking Universal Single-Copy Orthologs) is a popular tool used for this purpose, assessing the presence, fragmentation, or absence of highly conserved lineage-specific genes within an assembly (Simão et al. 2015; Manni et al. 2021). BUSCO first identifies candidate genomic regions using tBLASTn for protein homology searches (Camacho et al. 2009). These regions are then refined with AUGUSTUS (Stanke et al. 2004), a gene prediction tool trained on species-specific models, followed by alignment with HMMER (Eddy 2011) against curated ortholog databases. From version 5 onward, MetaEuk (Levy Karin et al. 2020) replaces the tBLASTn + AUGUSTUS pipeline for eukaryotic gene prediction and improving BUSCO’s scalability. However, the original pipeline is still required for prokaryotic and archaeal genome assessments, where MetaEuk is unsuitable. Based on its gene prediction analysis, BUSCO classifies genes into four categories: complete (single-copy) if found intact in a single copy, complete (duplicated) if multiple copies are detected, fragmented if only partial sequences are present, or missing if absent from the assembly. The BUSCO completeness score is the proportion of complete single-copy orthologs, providing a standardized metric by comparing the expected ortholog counts to those identified in the assembly.

Despite its widespread use, BUSCO has limitations. Its reliance on predefined ortholog sets can underestimate completeness for evolutionarily distant genomes (Gagalova et al. 2022). Furthermore, although the inclusion of MetaEuk improves its efficiency, BUSCO is still resource-intensive, requiring 30-80 minutes per assembly for large genomes, due to its dependence on sequence alignment and gene prediction algorithms (Huang and Li 2023).

Compleasm (Huang and Li 2023) is a faster alternative to BUSCO. It uses miniprot (Li 2023), a protein-to-genome alignment tool built on minimap2 (Li 2018), to accelerate conserved gene searches. By employing a minimizer-based (Roberts et al. 2004) mapping approach rather than traditional sequence alignment, compleasm achieves faster run times and lower memory usage while maintaining accuracy comparable to BUSCO without requiring gene prediction. Starting from BUSCO v5.7.0+, miniprot is used as the default search method, with the original tBLASTn + AUGUSTUS or MetaEuk pipeline still available as alternatives. Although these improvements reduce computational overhead (10-18 mins per assembly for large genomes), further run time reductions are needed as genome projects grow in size and complexity, underscoring the ongoing demand for faster completeness assessment methods.

Here we present aaKomp, an alignment-free genome completeness assessment tool that addresses these scalability challenges through amino acid *k-*mer matching. aaKomp employs aaHash (Wong et al. 2023b), a recursive hashing algorithm with BLOSUM62-based substitution tolerance (Henikoff and Henikoff 1992), combined with a multi-index Bloom filter (miBf) (Chu et al. 2020) for efficient *k*-mer storage and querying. This approach bypasses sequence alignment entirely while maintaining robust gene detection when there is sequence divergence between the analyzed genome and reference protein set.

Unlike existing tools, aaKomp computes a proportional completeness score that provides a finer resolution than threshold-based classifications and supports user-defined gene sets for customized assessments across any organism or lineage. Benchmarking on simulated and experimental datasets demonstrates that aaKomp maintains accuracy comparable to BUSCO and compleasm while achieving 68-fold and 18-fold faster execution, respectively, with up to 15-fold lower memory consumption. This improved performance makes aaKomp well-suited for large-scale genomic projects from population genomics to biodiversity studies.

## Methods

### 1. Algorithm

To evaluate genome completeness, aaKomp processes a target genome assembly against a reference protein set. This reference may be provided as raw sequences or as a pre-constructed multi-index Bloom filter (miBf) containing homology-aware amino acid *k*-mer hashes. The miBf functions as a probabilistic hash table that encodes metadata alongside *k-*mer presence. The miBf is constructed using the *k-*mer size of *k* (default: 9) and *h* (default: 9) number of hash functions, with a targeted false positive rate of *f* (default: 1%). Hashing is performed using aaHash, which supports three substitution-aware hashing levels: level 1 uses a strict 1:1 amino-acid mapping; level 2 merges residues with within-group BLOSUM62 substitution scores ≥1; and level 3 applies a more permissive merging of residues with within-group scores ≥0 (Tables S1–3).

The miBf includes a saturation detection feature that flags whether stored information may be ambiguous due to hash collisions. If all hash positions for a *k*-mer collide with existing entries, miBf marks the location as saturated to indicate that information may be missing or ambiguous. Conversely, an unsaturated mark ensures that no collision has occurred, and the encoded information is unambiguous.

To construct a custom miBf, each protein is first assigned a unique identifier and *k-*merized. Each *k-*mer is then associated with a 64-bit value: the first 32 bits represent the protein ID, and the second 32 bits store the *k-*mer’s position within the protein. This metadata is inserted into the miBf using the multi-level aaHash hash values of the *k-*mer. Additionally, for each protein, a targeted miBf is generated using a smaller *rescue_kmer* (default: 4) size to facilitate later recovery of short exonic regions.

aaKomp queries begin by performing a six-frame translation of the input genome assembly, independently processing both forward and reverse strands. For each strand, all three reading frames are analyzed to identify candidate seed regions (Fig. S1). A seed is defined as two consecutive aaHash Level 1 *k-*mers that share the same protein identifier and have monotonically increasing position values in the miBf. Once a seed is identified, extension is carried out only within the same frame where the seed is identified. Other frames advance concurrently, ensuring position consistency across all three reading frames.

During extension, all three substitution levels of aaHash are queried. The extension proceeds until the current *k-*mer’s hash values fail to satisfy the unsaturated status, protein ID and positional continuity constraints. The resulting segment is stored as a block, and the seed-extension process is repeated until each sequence has been processed.

The blocks are ordered sequentially in genomic coordinates and grouped by protein identifiers, and then chained together (Fig. S2). A lookahead of one block is performed to determine whether the next-next block results in a longer chain with a smaller positional gap. Block chaining continues until a non-viable block is encountered, when the start position of the next block is earlier than the end position of the current block.

If the total reconstruction, defined as the fraction of *k*-mers matching a specific protein ID that occur in monotonically increasing positions within evaluated blocks relative to the total number of *k*-mers in that protein, exceeds a threshold *l* (default: 70%), each gap between blocks is inspected and extracted. The targeted miBf with a reduced *k-*mer size is used to recover these missing regions. This recovery process searches for missing *k-*mers within the gap region and verifies that they appear in monotonically increasing order, without requiring seed identification. After recovery, the total number of *k*-mers across all the grouped blocks is summed. Because each exon boundary prevents the detection of *k* – 1 potential *k*-mers, this value is added to the total number of *k*-mers for the block for each putative exon boundary. The total value is then divided by the total number of *k-*mers in the protein sequence to compute the completeness score. A GFF3 annotation file is then generated for each evaluated protein stored in the miBf. Additionally, a Cumulative Distribution Function (CDF) is constructed using the best representation of each protein. The area under the curve (AUC) is calculated, and (1 − *AUC*) × 100 is reported as the final aaKomp score. Sequences shorter than *k* + 2 are disregarded in the calculation of the aaKomp score, as they cannot fulfill the initial seed requirement.

### 2. Implementation

aaKomp is implemented in C++ using btllib (v1.7.3) (Nikolić et al. 2022), libsequence (v1.9.8) (Thornton 2003), and boost (v1.85.0) libraries. The core algorithm and the ‘make_mibf’ module are compiled into standalone binaries. A lightweight Python driver script is included to facilitate command-line execution and ease of use. An optional R script is provided to generate CDF plots for visualization using dplyr) (Wickham et al. 2023) and ggplot2 (Wickham 2016).

### 3. Evaluation

To evaluate the performance of aaKomp (v1.0.0) against BUSCO (v5.8.3) and compleasm (v0.2.7), we used the T2T-CHM13 (v2.0) genome assembly (Accession: GCA_009914755.4) as a ground truth reference for generating genomes with simulated breakpoints (Table S4 and Method S1). Breakpoints were introduced in 2,000-interval steps, ranging from 0 to 98,000, to create assemblies with varying degrees of fragmentation and a wide range of contiguity levels (N50 range: 53.42 kbp–150.62 Mbp), reflecting the spectrum typically observed in real genome assemblies. A custom Python script was used to insert breakpoints at random positions in the reference-grade assembly (see the Code Availability section). Additionally, we obtained genome assemblies from the HPRC Year 1 freeze (https://github.com/human-pangenomics/HPP_Year1_Assemblies/blob/main/assembly_index/Year1_assemblies_v2_genbank.index) to assess performance on real-world data. For all tools, the ‘primates_odb12’ OrthoDB lineage was used to ensure consistency across comparisons (Tegenfeldt et al. 2025).

To evaluate aaKomp’s performance using a more closely related reference database, we used the human proteome from UniProt (UP000005640, https://www.uniprot.org/proteomes/UP000005640) (Ahmad et al. 2025). We assessed the T2T-CHM13 reference and the 94 HPRC genome assemblies using aaKomp with the human proteome to compare completeness scores and computational requirements against those obtained using the curated ortholog lineage.

To evaluate performance across diverse taxa, we assessed the *Anguilla anguilla* (European eel) genome assembly (GCA_013347855.1). Evaluation was performed using both the ‘actinopterygii_odb112 ortholog lineage and the complete *A. anguilla* proteome (UP001044222, https://www.uniprot.org/proteomes/UP001044222) (Ahmad et al. 2025).

To construct reference databases compatible with aaKomp, we used ‘hmmemit -c’ within HMMER (v3.1b2) (Eddy 2011) software suite to generate consensus sequences from the BUSCO ‘primates_odb12’ and ‘actinopterygii_odb12’ HMM ortholog profiles (Manni et al. 2021). These consensus sequences, along with the complete human and European eel proteomes, were used as input to the ‘make_mibf’ module of aaKomp.

We conducted a parameter sensitivity analysis by performing a *k*-mer sweep on the T2T-CHM13 assembly using the ‘primates_odb12’ miBf database. We tested *k*-mer sizes ranging from 7 to 11 to determine the impact of *k*-mer length on scoring accuracy.

All benchmarking tests were performed on a server-class system with 144 Intel(R) Xeon(R) Gold 6254 CPUs @ 3.1 GHz and 2.9 TB RAM using 48 threads.

## Results

### 1. Simulated dataset

To benchmark accuracy, we compared aaKomp against BUSCO and compleasm using a controlled ground truth dataset derived from the T2T-CHM13 [34] human genome. We systematically assessed correlations between the methods and measured their computational resource usage across genome assemblies with varying contiguity levels. For all comparisons, we used the ‘primates_odb12’ ortholog lineage for both BUSCO and compleasm, and generated a corresponding aaKomp miBf database from consensus sequences derived from the lineage’s HMM profiles. We compared aaKomp scores to BUSCO and compleasm completeness values, including both single-copy and duplicated genes.

aaKomp demonstrated high correlation with both BUSCO and compleasm, with Pearson correlation coefficients of *r* = 0.9995 and confidence intervals of [0.9991, 0.9997] and of *r* = 0.9996 and confidence intervals of [0.9993, 0.9998], respectively (Fig. 1) (Freedman et al. 2020). Across the full range of introduced breakpoints (0–98,000), aaKomp scores (78.55–94.03) remained closely aligned with the single-copy and duplicated completeness percentages reported by BUSCO (68.16%–100.00%) and compleasm (68.33%–100.00%). Using robust linear regression of aaKomp scores against respective tools’ completeness percentages, we observed slopes of 0.49 for BUSCO and 0.49 for compleasm, with regression lines fitting the majority of points (Liu et al. 2018).

**Figure 1.**
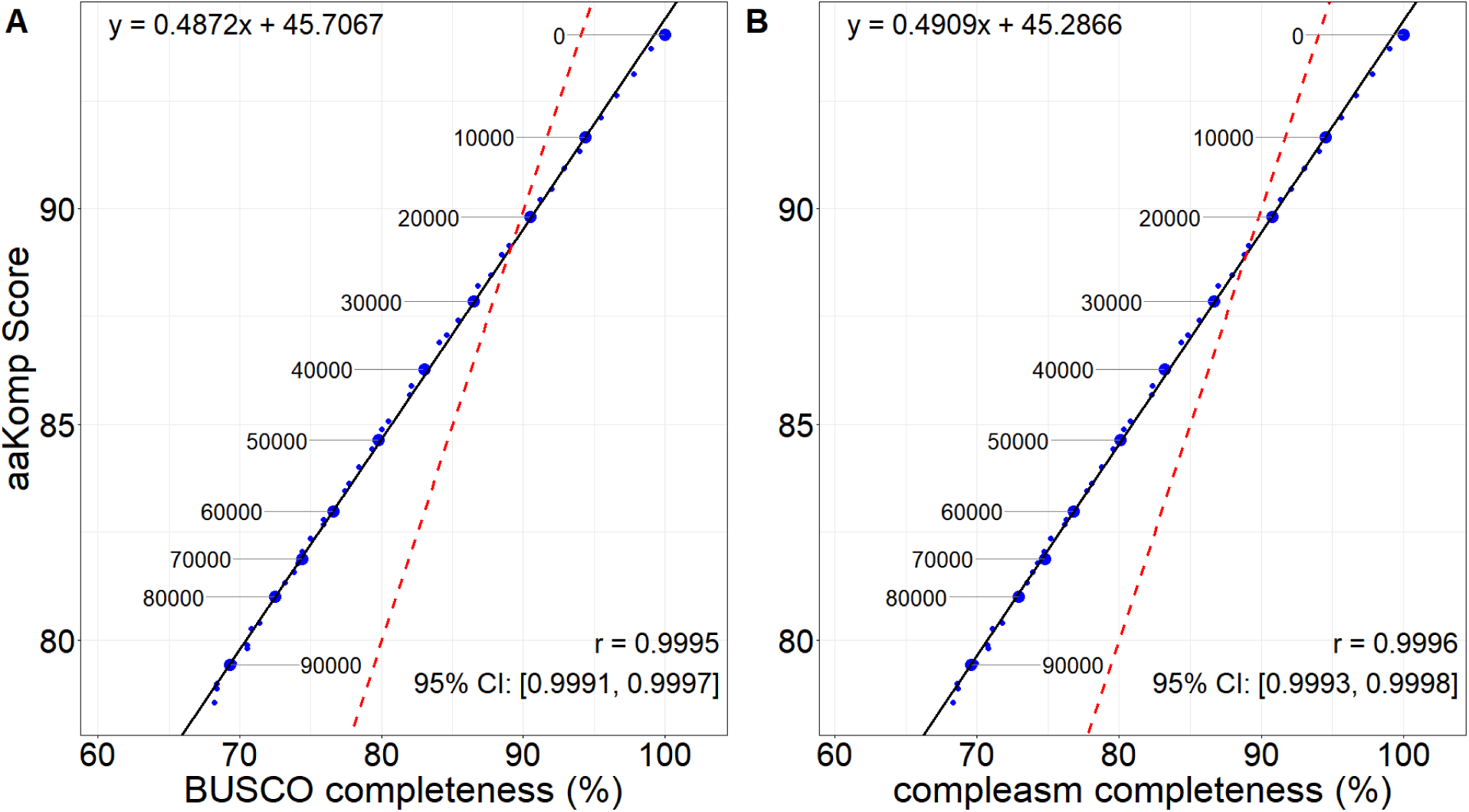
Correlation of gene set completeness scores between aaKomp and BUSCO, and aaKomp and compleasm, across simulated breakpoint assemblies of the CHM13-T2T v2.0 genome. **A:** Comparison of aaKomp scores with BUSCO gene completeness percentages across genome assemblies with varying numbers of simulated breakpoints (0–98,000). **B:** Comparison of aaKomp scores with compleasm gene completeness percentages for the same dataset. In both panels, individual points represent unique assemblies, with specific breakpoint intervals (0–90,000; 10,000-breakpoint step size) enlarged and annotated for clarity. The solid black line represents a robust linear regression fit, while the dashed red line represents the y = x line. Regression line equations are annotated at the top-left corner of each plot. *r* denotes the Pearson correlation coefficient, and CI the 95% confidence intervals.

For all tested genome assemblies, aaKomp outperformed BUSCO and compleasm in run time (Fig. 2A). It completed gene completeness assessments in an average of 0.58 ± 0.15 min (range: 0.55–1.61 min), approximately 68X faster on average than BUSCO (39.32 ± 10.78 min, range: 17.41–82.00 min) and 23X faster than compleasm (13.52 ± 0.79 min, range: 12.25–15.94 min). BUSCO exhibited the widest run time distribution, with several assemblies taking over an hour to complete, while compleasm was more consistent but still substantially slower than aaKomp. Across all 50 assemblies, aaKomp required a total of 29.13 min, compared to 1,965.94 min for BUSCO (∼32.77 h) and 675.86 min (∼11.26 h) for compleasm.

**Figure 2.**
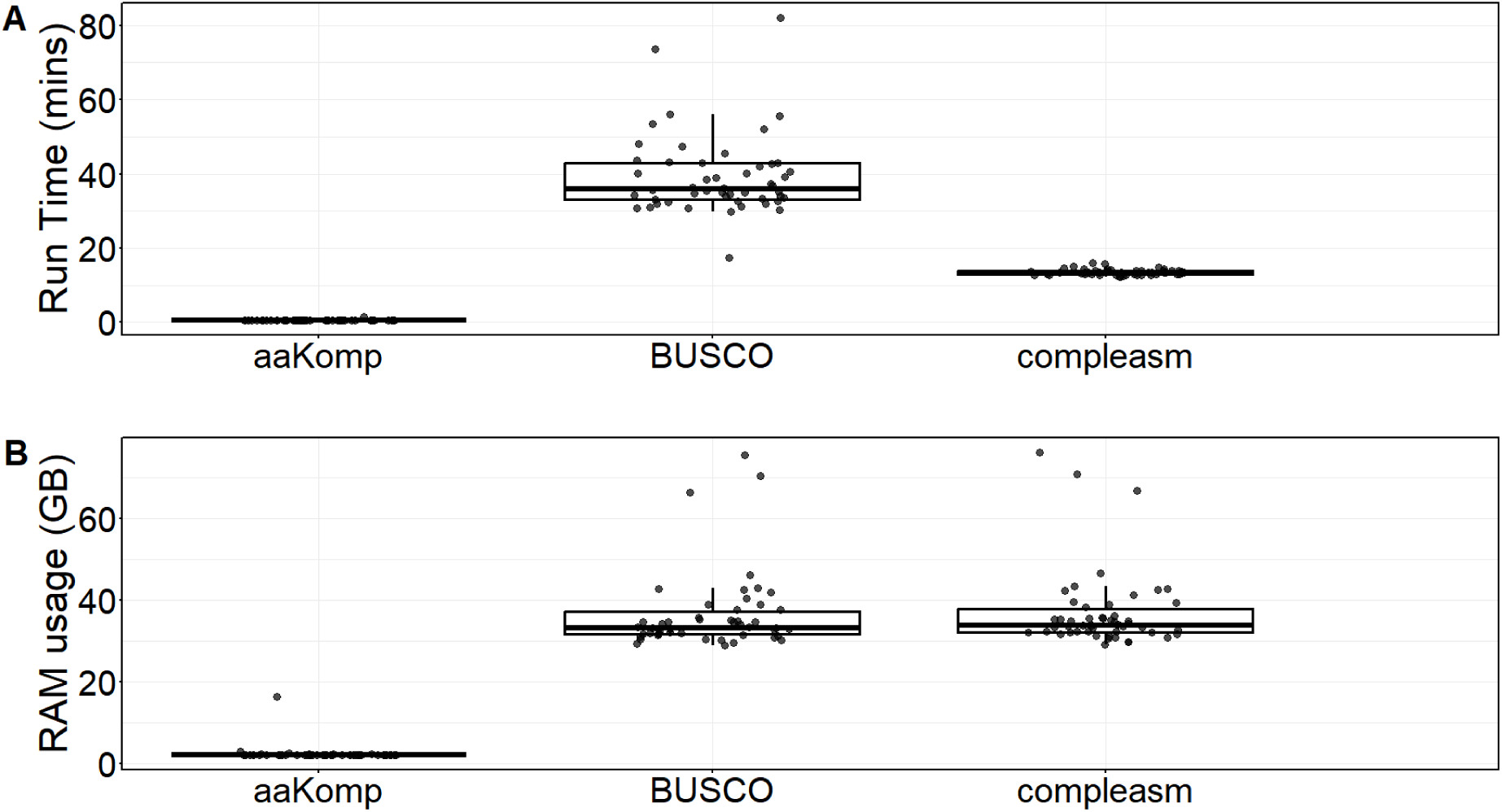
Run time and memory usage comparisons between aaKomp, BUSCO, and compleasm on simulated breakpoint assemblies of the CHM13-T2T v2.0 genome. **A:** Run time comparison for aaKomp, BUSCO, and compleasm across all simulated breakpoint assemblies of the CHM13-T2T v2.0 genome (0–98,000 breakpoints; n = 50). **B:** Memory usage (GB) comparison for the same set of assemblies. Each point represents an individual assembly run. Boxplots indicate the distribution of run times and memory consumption

Memory usage followed a similar pattern, with aaKomp requiring substantially less memory across all assemblies compared to both BUSCO and compleasm, with its most RAM-intensive run using roughly half as much memory (16.26 GB) as the least memory-intensive BUSCO or compleasm evaluation (Fig. 2B). aaKomp consumed an average of 2.44 ± 1.98 GB of RAM (range: 2.01–16.26 GB) across all assemblies. In contrast, BUSCO and compleasm required substantially more memory, with mean usages of 36.42 ± 9.63 GB (range: 28.91–75.52 GB) and 36.94 ± 9.61 GB (range: 29.19–76.03 GB), respectively, approximately 15X higher than aaKomp on average.

### 2. Human Pangenome Reference Consortium Genome Assemblies

We evaluated the performance of aaKomp across 94 genome assemblies from the HPRC Year 1 Freeze, including 47 maternal and 47 paternal haplotypes with the same miBf database used in testing the simulated dataset (Fig. 3). The aaKomp scores were consistent across genomes, with a score (aaKomp Score Mean ± S.D.) of 93.69 ± 0.91 and only a 2.82% difference between the lowest and highest scoring genomes (Fig. 3A). The scores formed two distinct clusters: the lower-scoring cluster corresponded to paternal haplotypes of male individuals, while the higher-scoring cluster included all other assemblies.

**Figure 3.**
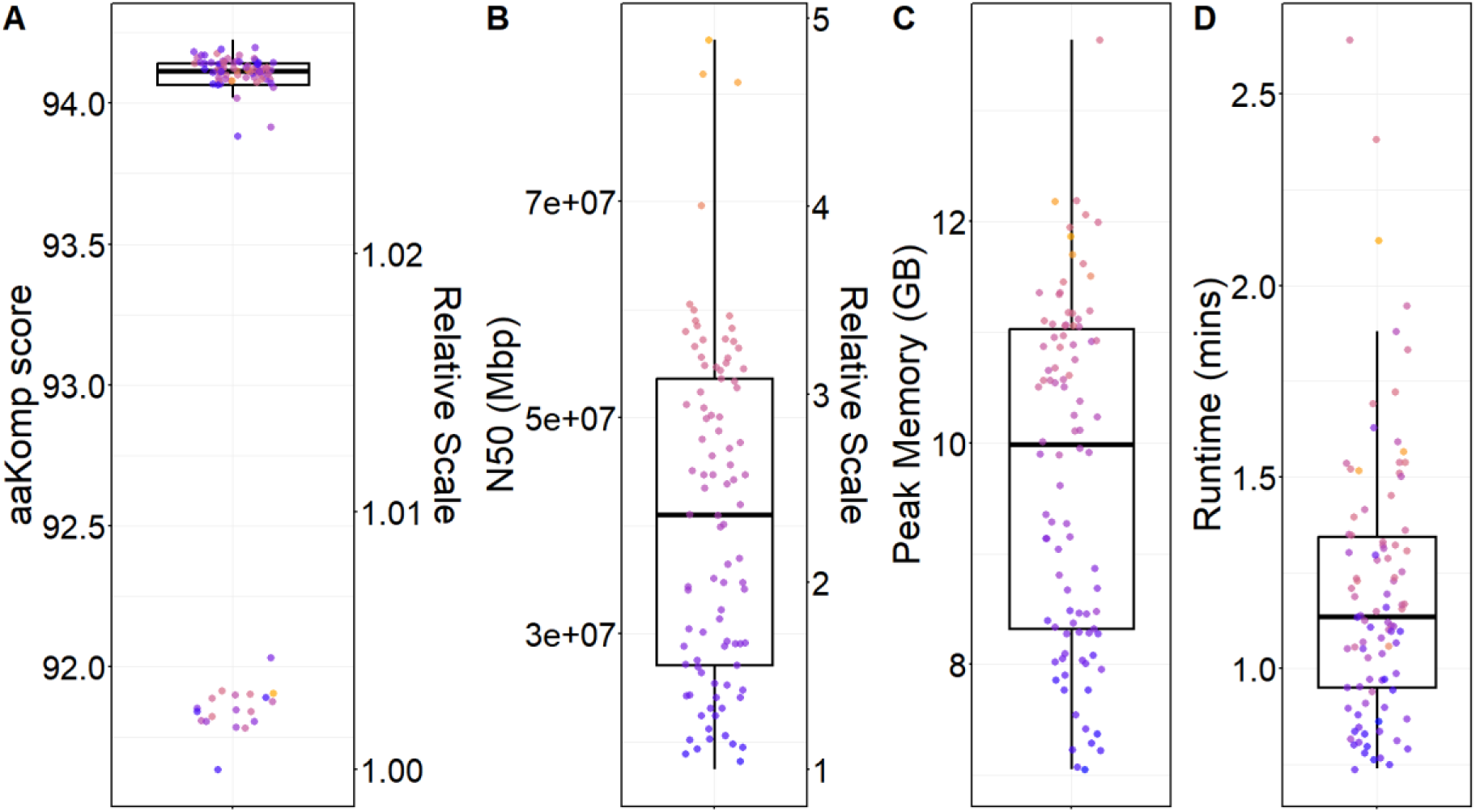
Performance of aaKomp across Human Pangenome Reference Consortium assemblies assessed using the ‘primates_odb12’ miBf database. **A:** aaKomp scores across Year 1 HPRC assemblies (n = 94), with the right axis showing the same values on a relative scale. **B:** Genome assembly contiguity (N50, in Mbp) with corresponding relative scale. **C:** Peak memory usage (GB) per aaKomp run. **D:** Run time (min) per aaKomp run. Each point represents an individual genome assembly. Boxplots indicate the distribution of values across all assemblies, with colour gradient representing the range of N50 values.

Assembly contiguity, measured by the N50 length metric—the length at which 50% of the genome is contained in contigs of that length or longer—ranged from 17.44 Mbp to 84.97 Mbp (Fig. 3B). aaKomp used less than 13.64 GB of memory across all runs, with most assemblies requiring (Peak RAM Mean ± S.D.) 9.74 ± 1.52 GB of RAM (Fig. 3C), and completed all assessments with a mean run time of 1.20 ± 0.35 min and a maximum of 2.64 min (Fig. 3D). Genome assemblies with higher contiguity tended to require more time and memory. The aaKomp score, however, showed greater variability across assemblies from different individuals, regardless of contiguity.

### 3. Running aaKomp using the human proteome

To assess aaKomp’s flexibility with user-defined databases and evaluate performance with a larger reference set, we repeated the HPRC analysis using a miBf database generated from the complete human proteome rather than the curated ‘primates_odb12’ ortholog set.

We evaluated aaKomp’s performance on the T2T-CHM13 reference genome and across 94 genome assemblies from the HPRC Year 1 Freeze, using a miBf database generated from the human proteome (UP000005640). On the T2T-CHM13 v2.0 genome assembly, aaKomp achieved a score of 97.48, with a peak memory usage of 22.77 GB and a run time of 5.45 min. Across the HPRC assemblies, aaKomp reported a mean score of 96.78 ± 1.03 with only a 3.05% difference between the lowest and highest assemblies, forming two distinct clusters (Fig. 4A). Memory usage remained below 19.00 GB for all runs, with most assemblies requiring 13.56 ± 2.05 GB of RAM (Fig. 4B). Run time averaged 2.96 ± 1.17 min, with the longest run completing in 7.33 min (Fig. 4C). A similar trend was observed as in the experiment with the ‘primates_odb12’ miBf database: run time and memory usage scaled with assembly contiguity, while aaKomp scores showed greater variability across individuals and did not strongly correlate with contiguity.

**Figure 4.**
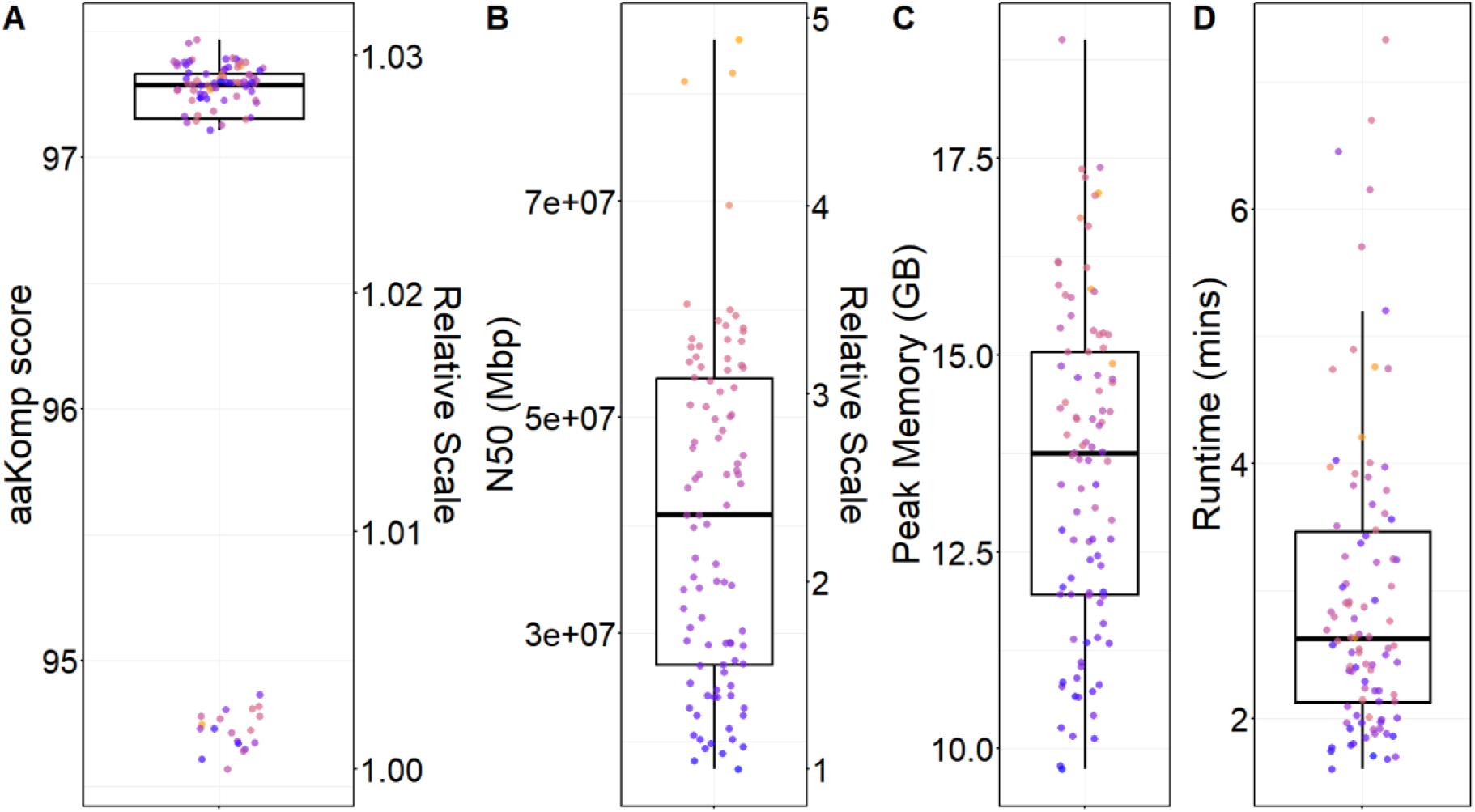
Performance of aaKomp across Human Pangenome Reference Consortium assemblies using the human proteome miBf database. **A:** aaKomp scores across Year 1 HPRC assemblies (n = 94), with the right axis showing the same values on a relative scale. **B:** Genome assembly contiguity (N50, in Mbp) with corresponding relative scale. **C:** Peak memory usage (GB) per aaKomp run. **D:** Run time (min) per aaKomp run. Each point represents an individual genome assembly. Boxplots indicate the distribution of values across all assemblies, with colour gradient representing the range of N50 values.

### 4. Performance on the European Eel (*Anguilla anguilla*) Genome

To evaluate aaKomp’s performance on a non-human vertebrate, we assessed the *A. anguilla* genome assembly. We performed evaluations using two distinct reference sets: the curated Actinopterygii (ray-finned fish) ortholog lineage (‘actinopterygii _odb12’) and the complete *A. anguilla* proteome.

Using the ‘actinopterygii_odb12’ miBf database, aaKomp achieved a score of 49.21, completing the analysis in 1.08 minutes with a peak memory of 5.55 GB. When evaluated against the European eel proteome, the aaKomp score was 94.79, requiring 1.75 minutes and 9.74 GB of RAM. Similar to the human benchmarks, the proteome-based analysis utilized a more closely related reference set, resulting in a higher aaKomp score.

### 5. Construction of miBf Databases

We also profiled the construction of the four distinct miBf reference databases used in the experiments. The construction of the ‘primates_odb12’ and human proteome miBf databases used in the human genome completeness assessments required 1.39 and 2.01 minutes, with peak memory usages of 1.97 and 3.15 GB, respectively. For the non-human benchmarks, we constructed miBf databases for the ‘actinopterygii_odb12’ lineage and the *A. anguilla* proteome, requiring 0.68 and 2.55 minutes, with peak memory usages of 1.18 and 3.53 GB, respectively.

### 6. Sensitivity to *k*-mer Size

We performed a parameter sweep to determine the impact of *k*-mer length on aaKomp’s sensitivity and computational performance. Across a tested *k* range from 7 to 11 using the T2T-CHM13 assembly and the ‘primates_odb12’ database, we found that a *k*-mer size of 9 yielded the highest aaKomp score of 94.03 (Table S5).

## Discussion

aaKomp utilizes an alignment-free approach to genome completeness assessment, employing fast, substitution-aware amino acid *k*-mer comparisons and supporting customizable reference inputs. Unlike traditional state-of-the-art tools that rely on computationally intensive sequence alignments, aaKomp evaluates completeness by quantifying the *k*-mer representation of reconstructed genes, capturing biologically meaningful variation through substitution tolerance. This is achieved by leveraging aaHash, a recursive hashing algorithm informed by BLOSUM62 substitution matrices, alongside an miBf, a data structure that efficiently stores *k*-mers with associated gene and positional metadata to enable rapid, homology-aware sequence queries (Henikoff and Henikoff 1992; Chu et al. 2020; Wong et al. 2023b).

This approach translates to high computational efficiency and strong empirical agreement with existing tools across all benchmarking tests. In terms of resource requirements, mean run times ranged from 1.20 to 2.96 minutes with average memory usage between 9.74 GB and 13.56 GB, scaling with the size of the reference set. In our simulation experiments, aaKomp saved over 10.78 hours and 32.28 hours of runtime compared to compleasm and BUSCO, respectively, across 50 assemblies. Despite this reduced overhead, we observed high score concordance with the alignment-based methods, achieving correlation coefficients of *r* = 0.9995 with BUSCO and *r* = 0.9996 with compleasm. aaKomp’s use of the miBf data structure also enables rapid custom database generation; for instance, custom databases for both the primates lineage and human proteome were generated in approximately 2 minutes. When applied to the T2T-CHM13 v2.0 genome assembly, the human proteome reference yielded a higher completeness score of 97.48, illustrating the gains of using a more proximal reference, whereas the use of the ‘primates_odb12’ lineage resulted in a score of 94.03, albeit with a shorter runtime and lower memory usage (S1 Discussion).

In larger projects involving hundreds or thousands of assemblies, the time savings observed in our results can potentially accelerate project timelines by weeks or even months. Such gains are made possible by an alignment-free design that enables faster and more scalable completeness assessments, making it well-suited for large-scale genome projects like the Earth BioGenome Project (EBP) and HPRC where researchers might be interested in monitoring results across every step of multi-stage protocols, or assembly strategies involve optimizing a combinatorial number of genome assembly parameters, such as GoldRush (Lewin et al. 2018; Liao et al. 2023; Wong et al. 2023a; “HPRC Assembly Production” 2024; “Report on Assembly Recommendations” 2026; “Report on Assembly Standards” 2026). This time efficiency and lower memory usage are achieved in part by the miBf, which builds on the Bloom filter—a space-efficient probabilistic data structure designed to test set membership—and incorporates an interleaved rank array with a dense value array (Chu et al. 2020). Unlike a standard Bloom filter, the miBf supports key-value association to reduce false positives through value verification. We further reduce these errors by encoding positional metadata—gene ID and *k*-mer position—into each value, providing a secondary layer of validation during the querying process.

To query the miBf and assess gene content, aaKomp uses aaHash, a recursive amino acid *k*-mer hashing algorithm that avoids the computational cost of traditional dynamic programming by leveraging multi-level hashing (Wong et al. 2023b). Instead of evaluating all possible alignments with substitutions, aaHash computes three hash representations for each *k*-mer across substitution-tolerant levels based on BLOSUM62, which help aaKomp detect conservative amino acid substitutions during matching. The substitution-tolerant hashing enables aaKomp to recover and score genes even when the assembled sequence does not perfectly match the reference, extending its utility to organisms whose genomes harbor some degree of sequence divergence from the reference gene set. aaKomp leverages the rolling hash mechanism of aaHash to efficiently query translated sequences, computing each new *k*-mer hash by incrementally updating the previous value rather than rehashing from scratch. This combination of recursive and homology-aware hashing and a space-efficient index allows aaKomp to evaluate gene content in genomic data an order of magnitude faster than state-of-the-art tools with a minimal RAM footprint.

While the computational efficiency of aaKomp is a significant advantage, the tool’s utility ultimately depends on its ability to accurately reconstruct gene models across diverse biological contexts. Our results demonstrate high concordance with traditional methods, which is achieved through a combination of *k*-mer rescue strategies and strict positional constraints. Specifically, *k*-mers contributing to a gene’s completeness must originate from the same protein and occur at monotonically increasing positions, incrementing by one for each subsequent *k*-mer. This positional requirement ensures that only biologically consistent and contiguous matches are retained during evaluation. By utilizing the gene identifiers and coordinates associated with each *k*-mer, aaKomp is able to verify each gene product’s structural coherence. However, in protein sequence datasets with many repeated *k*-mers, frequent hash collisions can overwrite this information, preventing aaKomp from reconstructing the protein. To mitigate this loss, the miBf incorporates a saturation detection mechanism that identifies entries where all corresponding hash positions have suffered collisions. In these instances, aaKomp validates saturated *k*-mers lacking direct positional metadata by anchoring them to preceding unsaturated *k*-mers that carry consistent gene and coordinate information. This mechanism allows for the recovery of valid matches despite localized ambiguity, such as those found in common, widely distributed protein domains.

To ensure the recovery of sequences shorter than *k* during gene reconstruction, aaKomp must address the inherent limitations of single *k* approaches. Specifically, using a fixed *k*-mer size prevents the detection of sequences shorter than *k*, which leads to the omission of short exons and thus incomplete completeness assessments. To supplement its primary reconstruction framework, aaKomp employs a dual-layered approach: a general miBf (larger *k*-mer) to identify confident gene segments within the genome and a targeted miBf (smaller *k*-mer) to recover matches within these short exonic regions. This hybrid indexing strategy ensures a more robust assessment of genomic completeness by capturing the small structural details that larger *k*-mers would otherwise overlook. Our experiments sweeping on *k*-mer lengths indicated that a default length of nine with a rescue length of four yielded the highest aaKomp score, representing the optimal balance between detection accuracy and computational noise (Table S5).

aaKomp was designed to handle frameshift events by searching alternative reading frames on the same strand whenever *k*-mer matching is disrupted. Methodologically, aaKomp treats mismatches, insertions, deletions, and splice junctions as equivalent interruptions to the expected *k*-mer pattern. When a disruption is detected, aaKomp attempts to recover *k-*mers by searching all three reading frames, while simultaneously penalizing *k*-mer absences, such as those caused by deletions or non-conservative substitutions. This strategy enables aaKomp to distinguish disrupted gene regions from true genic loss.

Finally, aaKomp does not initiate a targeted miBf recovery search unless a significant fraction—by default, at least 70%—of the gene has already been reconstructed in the primary, more restrictive *k*-mer space. This threshold reduces the risk of spurious recovery from low-confidence regions while still allowing for the detection of genes containing short exons. Collectively, these strategies ensure that aaKomp’s completeness estimates remain both accurate and robust across diverse biological contexts.

Although aaKomp produces scores that closely correlate with existing methods, it takes a different approach to quantifying gene completeness. BUSCO and compleasm apply hard and arbitrary thresholds (80% of aligned gene length) to classify genes as either complete, fragmented or missing, whereas aaKomp quantifies gene recovery proportionally in *k-*mer space, capturing the degree of reconstruction. Threshold-based classifications may obscure reconstruction gaps—a gene at 80% recovery is classified identically to one at 100%, masking the true degree of completeness—and complicate interpretation, as during assembly parameter optimizations, one assembly could exhibit worse gene recovery yet appear equivalent in completeness score to another. In contrast, aaKomp’s proportional scoring produces a continuum of gene recovery across the entire completeness spectrum, making it a more nuanced and interpretable metric for tracking incremental improvements in genome assembly quality.

This also means that, in practice, aaKomp will rarely, if ever, report a completeness score of 100, as achieving this requires the presence of virtually all expected *k*-mers without exception. This is particularly relevant when evaluating with consensus sequences like that of OrthoDB lineages, where evolutionary divergence, natural polymorphisms, or incomplete reconstruction of short or variable exons may result in the absence of expected *k*-mers (Tegenfeldt et al. 2025). Even disregarding assembly artifacts or imperfections, genuine biological variation alone can introduce enough sequence differences to preclude perfect recovery in the *k*-mer space. Any substitution not tolerated by aaHash’s BLOSUM62-based scheme or even just a single amino acid deletion can prevent a perfect score from being achieved (Henikoff and Henikoff 1992). Rather than masking these gaps with threshold-based categories, aaKomp’s proportional scoring reflects them transparently. As such, individual scores are best interpreted comparatively, highlighting relative differences in gene recovery across assemblies rather than as absolute measures of completeness. By extension, aaKomp performs optimally when the reference database is evolutionarily close to the target genome, ensuring minimal *k*-mer divergence. As reflected in our benchmarking results, leveraging a more closely related proteome significantly enhances recovery, highlighting the importance of reference selection in minimizing *k*-mer mismatch.

BUSCO and compleasm also rely on predefined ortholog sets, which can limit their applicability when working with species that only have evolutionarily distant lineage references available. In such cases, researchers may be forced to use references that do not fully reflect the gene content of the target organism, simply due to the absence of closer alternatives. Indeed, the EBP genome assessment guidelines acknowledge that completeness targets may be relaxed for taxa strongly divergent from established BUSCO sets (“Report on Assembly Standards” 2026). In contrast, aaKomp allows users to generate customized reference databases, enabling lineage- or clade-specific assessments, by providing the proteome FASTA file, as demonstrated in the human and European eel proteome experiments. Notably, even with reference miBf database construction included, aaKomp still completed assessments faster than BUSCO or compleasm in the human experiments.

By eliminating the dependency on fixed ortholog sets and supporting rapid, user-defined reference database generation, aaKomp offers a highly flexible and adaptable solution across a wide range of taxa. Its speed and scalability enable researchers to efficiently evaluate and compare large numbers of assemblies, facilitating the detailed tracking of incremental improvements across assembly versions and addressing limitations of current state-of-the-art tools. Collectively, these advantages position aaKomp as a practical framework for modern large-scale genome projects.

## Disclaimer

Generative AI was used to assist in proofreading and refining the language of this article, and debugging the code. All scientific content, analysis, and conclusions are the sole work of the authors.

## Author contributions

J.W. and R.L.W. conceived the research project. J.W. implemented the code, performed analysis, and wrote the manuscript. L.C., R.L.W, and I.B. provided valuable discussion and suggestions. R.L.W and I.B. suggested edits to the manuscript and supervised the entire research.

## Code Availability

aaKomp (v1.0.0) has been deposited in Zenodo at https://doi.org/10.5281/zenodo.19124729. aaKomp and custom scripts used in the experiments are available at https://github.com/BirolLab/aakomp and released under the GPL-3 license.

## Data Availability

The ‘primates_odb12’, human proteome, ‘actinopterygii _odb12’, and European eel proteome miBf databases generated in this study have been deposited in Zenodo at https://doi.org/10.5281/zenodo.15421505, https://doi.org/10.5281/zenodo.15732544, https://doi.org/10.5281/zenodo.19124501, and https://doi.org/10.5281/zenodo.19124550 respectively. The raw amino acid sequences used in building these databases are included with their respective miBfs. The breakpoint-fragmented genome assemblies generated for the simulated experiments described in Fig.1 are available from the authors upon request.

## References

Ahmad, S., Jose da Costa Gonzales, L., Bowler-Barnett, E.H., Rice, D.L., Kim, M., Wijerathne, S., Luciani, A., Kandasaamy, S., Luo, J., Watkins, X., Turner, E., Martin, M.J., and the UniProt Consortium. 2025. The UniProt website API: facilitating programmatic access to protein knowledge. Nucleic Acids Res 53(W1): W547–W553. doi:10.1093/nar/gkaf394.

Camacho, C., Coulouris, G., Avagyan, V., Ma, N., Papadopoulos, J., Bealer, K., and Madden, T.L. 2009. BLAST+: architecture and applications. BMC Bioinformatics 10(1): 421. doi:10.1186/1471-2105-10-421.

Cheng, H., Concepcion, G.T., Feng, X., Zhang, H., and Li, H. 2021. Haplotype-resolved de novo assembly using phased assembly graphs with hifiasm. Nat Methods 18(2): 170–175. Nature Publishing Group. doi:10.1038/s41592-020-01056-5.

Chu, J., Mohamadi, H., Erhan, E., Tse, J., Chiu, R., Yeo, S., and Birol, I. 2020. Mismatch-tolerant, alignment-free sequence classification using multiple spaced seeds and multiindex Bloom filters. Proceedings of the National Academy of Sciences 117(29): 16961–16968. Proceedings of the National Academy of Sciences. doi:10.1073/pnas.1903436117.

Eddy, S.R. 2011. Accelerated Profile HMM Searches. PLOS Computational Biology 7(10): e1002195. Public Library of Science. doi:10.1371/journal.pcbi.1002195.

Freedman, D., Pisani, R., and Purves, R. 2020. Statistics: Fourth international student edition. W.W. Norton & Company 22.

Gagalova, K.K., Warren, R.L., Coombe, L., Wong, J., Nip, K.M., Yuen, M.M.S., Whitehill, J.G.A., Celedon, J.M., Ritland, C., Taylor, G.A., Cheng, D., Plettner, P., Hammond, S.A., Mohamadi, H., Zhao, Y., Moore, R.A., Mungall, A.J., Boyle, B., Laroche, J., Cottrell, J., Mackay, J.J., Lamothe, M., Gérardi, S., Isabel, N., Pavy, N., Jones, S.J.M., Bohlmann, J., Bousquet, J., and Birol, I. 2022. Spruce giga-genomes: structurally similar yet distinctive with differentially expanding gene families and rapidly evolving genes. The Plant Journal 111(5): 1469–1485. doi:10.1111/tpj.15889.

Henikoff, S., and Henikoff, J.G. 1992. Amino acid substitution matrices from protein blocks. Proceedings of the National Academy of Sciences 89(22): 10915–10919. Proceedings of the National Academy of Sciences. doi:10.1073/pnas.89.22.10915.

HPRC Assembly Production. 2024, June 5. Available from https://github.com/human-pangenomics/hpp_production_workflows/tree/master/assembly.

Huang, N., and Li, H. 2023. compleasm: a faster and more accurate reimplementation of BUSCO. Bioinformatics 39(10): btad595. doi:10.1093/bioinformatics/btad595.

Jackman, S.D., Vandervalk, B.P., Mohamadi, H., Chu, J., Yeo, S., Hammond, S.A., Jahesh, G., Khan, H., Coombe, L., Warren, R.L., and Birol, I. 2017. ABySS 2.0: resource-efficient assembly of large genomes using a Bloom filter. Genome Res 27(5): 768–777. doi:10.1101/gr.214346.116.

Kolmogorov, M., Yuan, J., Lin, Y., and Pevzner, P.A. 2019. Assembly of long, error-prone reads using repeat graphs. Nat Biotechnol 37(5): 540–546. Nature Publishing Group. doi:10.1038/s41587-019-0072-8.

Levy Karin, E., Mirdita, M., and Söding, J. 2020. MetaEuk—sensitive, high-throughput gene discovery, and annotation for large-scale eukaryotic metagenomics. Microbiome 8(1): 48. doi:10.1186/s40168-020-00808-x.

Lewin, H.A., Robinson, G.E., Kress, W.J., Baker, W.J., Coddington, J., Crandall, K.A., Durbin, R., Edwards, S.V., Forest, F., Gilbert, M.T.P., Goldstein, M.M., Grigoriev, I.V., Hackett, K.J., Haussler, D., Jarvis, E.D., Johnson, W.E., Patrinos, A., Richards, S., Castilla-Rubio, J.C., van Sluys, M.-A., Soltis, P.S., Xu, X., Yang, H., and Zhang, G. 2018. Earth BioGenome Project: Sequencing life for the future of life. Proceedings of the National Academy of Sciences 115(17): 4325–4333. Proceedings of the National Academy of Sciences. doi:10.1073/pnas.1720115115.

Li, H. 2018. Minimap2: pairwise alignment for nucleotide sequences. Bioinformatics 34(18): 3094–3100. doi:10.1093/bioinformatics/bty191.

Li, H. 2023. Protein-to-genome alignment with miniprot. Bioinformatics 39(1): btad014. doi:10.1093/bioinformatics/btad014.

Li, H., and Durbin, R. 2024. Genome assembly in the telomere-to-telomere era. Nat Rev Genet 25(9): 658–670. Nature Publishing Group. doi:10.1038/s41576-024-00718-w.

Liao, W.-W., Asri, M., Ebler, J., Doerr, D., Haukness, M., Hickey, G., Lu, S., Lucas, J.K., Monlong, J., Abel, H.J., Buonaiuto, S., Chang, X.H., Cheng, H., Chu, J., Colonna, V., Eizenga, J.M., Feng, X., Fischer, C., Fulton, R.S., Garg, S., Groza, C., Guarracino, A., Harvey, W.T., Heumos, S., Howe, K., Jain, M., Lu, T.-Y., Markello, C., Martin, F.J., Mitchell, M.W., Munson, K.M., Mwaniki, M.N., Novak, A.M., Olsen, H.E., Pesout, T., Porubsky, D., Prins, P., Sibbesen, J.A., Sirén, J., Tomlinson, C., Villani, F., Vollger, M.R., Antonacci-Fulton, L.L., Baid, G., Baker, C.A., Belyaeva, A., Billis, K., Carroll, A., Chang, P.-C., Cody, S., Cook, D.E., Cook-Deegan, R.M., Cornejo, O.E., Diekhans, M., Ebert, P., Fairley, S., Fedrigo, O., Felsenfeld, A.L., Formenti, G., Frankish, A., Gao, Y., Garrison, N.A., Giron, C.G., Green, R.E., Haggerty, L., Hoekzema, K., Hourlier, T., Ji, H.P., Kenny, E.E., Koenig, B.A., Kolesnikov, A., Korbel, J.O., Kordosky, J., Koren, S., Lee, H., Lewis, A.P., Magalhães, H., Marco-Sola, S., Marijon, P., McCartney, A., McDaniel, J., Mountcastle, J., Nattestad, M., Nurk, S., Olson, N.D., Popejoy, A.B., Puiu, D., Rautiainen, M., Regier, A.A., Rhie, A., Sacco, S., Sanders, A.D., Schneider, V.A., Schultz, B.I., Shafin, K., Smith, M.W., Sofia, H.J., Abou Tayoun, A.N., Thibaud-Nissen, F., Tricomi, F.F., Wagner, J., Walenz, B., Wood, J.M.D., Zimin, A.V., Bourque, G., Chaisson, M.J.P., Flicek, P., Phillippy, A.M., Zook, J.M., Eichler, E.E., Haussler, D., Wang, T., Jarvis, E.D., Miga, K.H., Garrison, E., Marschall, T., Hall, I.M., Li, H., and Paten, B. 2023. A draft human pangenome reference. Nature 617(7960): 312–324. Nature Publishing Group. doi:10.1038/s41586-023-05896-x.

Liu, J., Cosman, P.C., and Rao, B.D. 2018. Robust Linear Regression via \ell_0 Regularization. IEEE Transactions on Signal Processing 66(3): 698–713. doi:10.1109/TSP.2017.2771720.

Manni, M., Berkeley, M.R., Seppey, M., Simão, F.A., and Zdobnov, E.M. 2021. BUSCO Update: Novel and Streamlined Workflows along with Broader and Deeper Phylogenetic Coverage for Scoring of Eukaryotic, Prokaryotic, and Viral Genomes. Molecular Biology and Evolution 38(10): 4647–4654. doi:10.1093/molbev/msab199.

Mikheenko, A., Prjibelski, A., Saveliev, V., Antipov, D., and Gurevich, A. 2018. Versatile genome assembly evaluation with QUAST-LG. Bioinformatics 34(13): i142–i150. doi:10.1093/bioinformatics/bty266.

Nikolić, V., Kazemi, P., Coombe, L., Wong, J., Afshinfard, A., Chu, J., Warren, R.L., and Birol, I. 2022. btllib: A C++ library with Python interface for efficient genomic sequence processing. Journal of Open Source Software 7(79): 4720. doi:10.21105/joss.04720.

Report on Assembly Recommendations. 2026, January. Available from https://www.earthbiogenome.org/report-on-assembly-recommendations.

Report on Assembly Standards. 2026, January. Available from https://www.earthbiogenome.org/report-on-assembly-standards-v7.

Rhie, A., McCarthy, S.A., Fedrigo, O., Damas, J., Formenti, G., Koren, S., Uliano-Silva, M., Chow, W., Fungtammasan, A., Kim, J., Lee, C., Ko, B.J., Chaisson, M., Gedman, G.L., Cantin, L.J., Thibaud-Nissen, F., Haggerty, L., Bista, I., Smith, M., Haase, B., Mountcastle, J., Winkler, S., Paez, S., Howard, J., Vernes, S.C., Lama, T.M., Grutzner, F., Warren, W.C., Balakrishnan, C.N., Burt, D., George, J.M., Biegler, M.T., Iorns, D., Digby, A., Eason, D., Robertson, B., Edwards, T., Wilkinson, M., Turner, G., Meyer, A., Kautt, A.F., Franchini, P., Detrich, H.W., Svardal, H., Wagner, M., Naylor, G.J.P., Pippel, M., Malinsky, M., Mooney, M., Simbirsky, M., Hannigan, B.T., Pesout, T., Houck, M., Misuraca, A., Kingan, S.B., Hall, R., Kronenberg, Z., Sović, I., Dunn, C., Ning, Z., Hastie, A., Lee, J., Selvaraj, S., Green, R.E., Putnam, N.H., Gut, I., Ghurye, J., Garrison, E., Sims, Y., Collins, J., Pelan, S., Torrance, J., Tracey, A., Wood, J., Dagnew, R.E., Guan, D., London, S.E., Clayton, D.F., Mello, C.V., Friedrich, S.R., Lovell, P.V., Osipova, E., Al-Ajli, F.O., Secomandi, S., Kim, H., Theofanopoulou, C., Hiller, M., Zhou, Y., Harris, R.S., Makova, K.D., Medvedev, P., Hoffman, J., Masterson, P., Clark, K., Martin, F., Howe, K., Flicek, P., Walenz, B.P., Kwak, W., Clawson, H., Diekhans, M., Nassar, L., Paten, B., Kraus, R.H.S., Crawford, A.J., Gilbert, M.T.P., Zhang, G., Venkatesh, B., Murphy, R.W., Koepfli, K.-P., Shapiro, B., Johnson, W.E., Di Palma, F., Marques-Bonet, T., Teeling, E.C., Warnow, T., Graves, J.M., Ryder, O.A., Haussler, D., O’Brien, S.J., Korlach, J., Lewin, H.A., Howe, K., Myers, E.W., Durbin, R., Phillippy, A.M., and Jarvis, E.D. 2021. Towards complete and error-free genome assemblies of all vertebrate species. Nature 592(7856): 737–746. Nature Publishing Group. doi:10.1038/s41586-021-03451-0.

Rhie, A., Walenz, B.P., Koren, S., and Phillippy, A.M. 2020. Merqury: reference-free quality, completeness, and phasing assessment for genome assemblies. Genome Biology 21(1): 245. doi:10.1186/s13059-020-02134-9.

Roberts, M., Hayes, W., Hunt, B.R., Mount, S.M., and Yorke, J.A. 2004. Reducing storage requirements for biological sequence comparison. Bioinformatics 20(18): 3363–3369. doi:10.1093/bioinformatics/bth408.

Satam, H., Joshi, K., Mangrolia, U., Waghoo, S., Zaidi, G., Rawool, S., Thakare, R.P., Banday, S., Mishra, A.K., Das, G., and Malonia, S.K. 2023. Next-Generation Sequencing Technology: Current Trends and Advancements. Biology 12(7): 997. Multidisciplinary Digital Publishing Institute. doi:10.3390/biology12070997.

Sayers, E.W., Beck, J., Bolton, E.E., Brister, J.R., Chan, J., Connor, R., Feldgarden, M., Fine, A.M., Funk, K., Hoffman, J., Kannan, S., Kelly, C., Klimke, W., Kim, S., Lathrop, S., Marchler-Bauer, A., Murphy, T.D., O’Sullivan, C., Schmieder, E., Skripchenko, Y., Stine, A., Thibaud-Nissen, F., Wang, J., Ye, J., Zellers, E., Schneider, V.A., and Pruitt, K.D. 2025. Database resources of the National Center for Biotechnology Information in 2025. Nucleic Acids Research 53(D1): D20–D29. doi:10.1093/nar/gkae979.

Shafin, K., Pesout, T., Lorig-Roach, R., Haukness, M., Olsen, H.E., Bosworth, C., Armstrong, J., Tigyi, K., Maurer, N., Koren, S., Sedlazeck, F.J., Marschall, T., Mayes, S., Costa, V., Zook, J.M., Liu, K.J., Kilburn, D., Sorensen, M., Munson, K.M., Vollger, M.R., Monlong, J., Garrison, E., Eichler, E.E., Salama, S., Haussler, D., Green, R.E., Akeson, M., Phillippy, A., Miga, K.H., Carnevali, P., Jain, M., and Paten, B. 2020. Nanopore sequencing and the Shasta toolkit enable efficient de novo assembly of eleven human genomes. Nat Biotechnol 38(9): 1044–1053. Nature Publishing Group. doi:10.1038/s41587-020-0503-6.

Simão, F.A., Waterhouse, R.M., Ioannidis, P., Kriventseva, E.V., and Zdobnov, E.M. 2015. BUSCO: assessing genome assembly and annotation completeness with single-copy orthologs. Bioinformatics 31(19): 3210–3212. doi:10.1093/bioinformatics/btv351.

Stanke, M., Steinkamp, R., Waack, S., and Morgenstern, B. 2004. AUGUSTUS: a web server for gene finding in eukaryotes. Nucleic Acids Res 32(Web Server issue): W309–W312. doi:10.1093/nar/gkh379.

Tegenfeldt, F., Kuznetsov, D., Manni, M., Berkeley, M., Zdobnov, E.M., and Kriventseva, E.V. 2025. OrthoDB and BUSCO update: annotation of orthologs with wider sampling of genomes. Nucleic Acids Res 53(D1): D516–D522. doi:10.1093/nar/gkae987.

Thornton, K. 2003. libsequence: a C++ class library for evolutionary genetic analysis. Bioinformatics 19(17): 2325–2327. doi:10.1093/bioinformatics/btg316.

Wickham, H. 2016. ggplot2: Elegant Graphics for Data Analysis. Springer-Verlag New York. Available from https://ggplot2.tidyverse.org.

Wickham, H., François, R., Henry, L., Müller, K., and Vaughan, D. 2023. dplyr: A Grammar of Data Manipulation. Available from https://dplyr.tidyverse.org.

Wong, J., Coombe, L., Nikolić, V., Zhang, E., Nip, K.M., Sidhu, P., Warren, R.L., and Birol, I. 2023a. Linear time complexity de novo long read genome assembly with GoldRush. Nat Commun 14(1): 2906. Nature Publishing Group. doi:10.1038/s41467-023-38716-x.

Wong, J., Kazemi, P., Coombe, L., Warren, R.L., and Birol, I. 2023b. aaHash: recursive amino acid sequence hashing. Bioinformatics Advances 3(1): vbad162. doi:10.1093/bioadv/vbad162.

zyndagj/ncbi_genomes. (n.d.). Available from https://github.com/zyndagj/ncbi_genomes [accessed 19 April 2025].

