## Supplementary Materials for "aaKomp: Alignment-free amino acid *k*-mer matching for genome completeness assessment at scale"

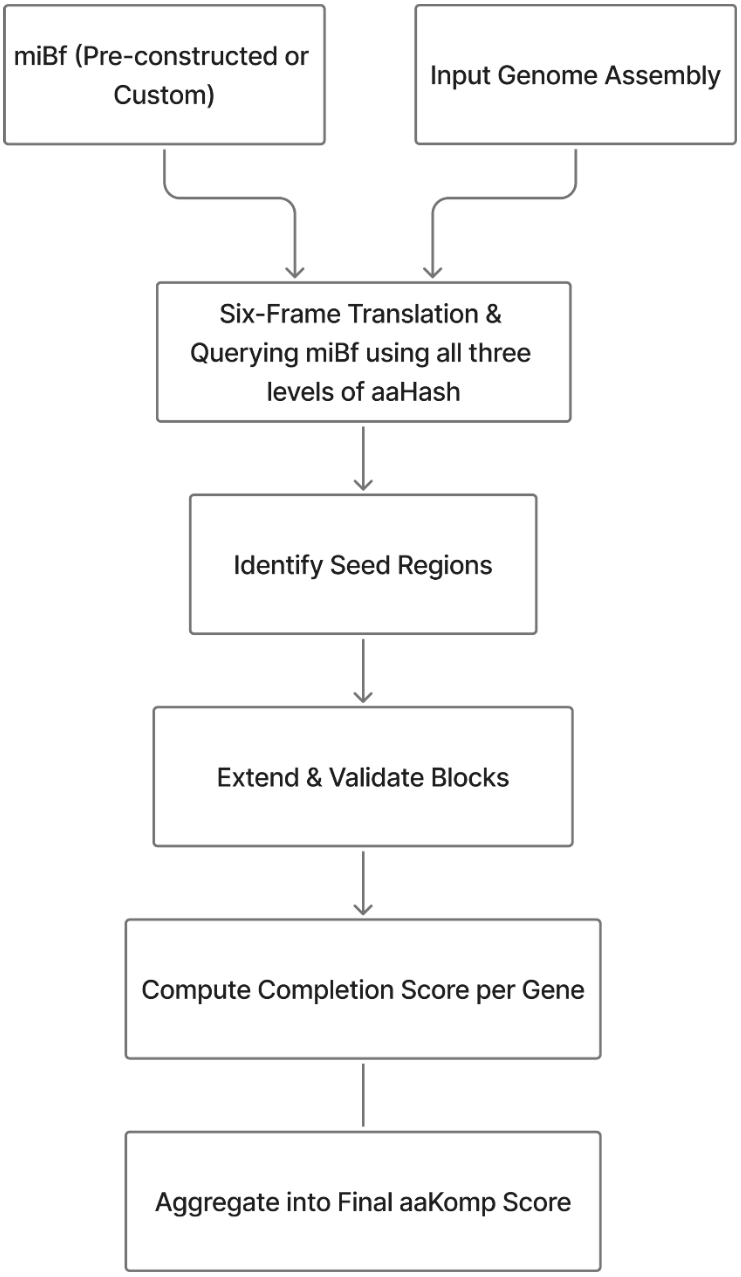

**Figure S1. High-level overview of the aaKomp workflow.** aaKomp will evaluate genic completeness by scanning an input genome assembly for homology-aware amino acid *k*-mer hashes stored in miBf, which may be pre-constructed or generated from a custom reference dataset. The assembly will be translated in all six reading frames and queried against the miBf using all three levels of aaHash. Seed regions will first be identified and subsequently extended and validated into contiguous blocks using consecutive *k*-mer hits. These blocks will then be used to compute a completion score for each gene, and the per-gene scores will be aggregated to form the final aaKomp score

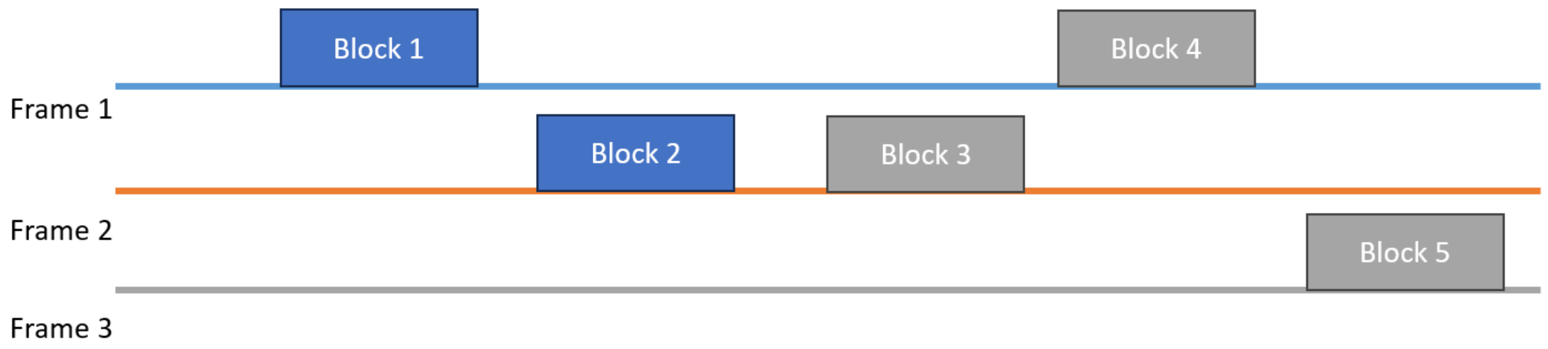

**Figure S2. Chaining of gene reconstruction blocks across different protein identifiers.** Blocks of monotonically increasing matched *k*-mers are first ordered by genomic coordinates, grouped by specific protein identifiers, and then chained together for downstream evaluation. In this example, the blue blocks (Blocks 1 and 2) share a single protein identifier, while the grey blocks (Blocks 3, 4, and 5) belong to another. Blocks with shared identifiers are chained together.

**Table S1. aaHash Level 1 hashing scheme**

$$\Sigma_1 = \{A, R, N, D, C, E, Q, G, H, I, L, K, M, F, P, S, T, W, Y, V\}$$

Defines the aaHash hashing scheme in which each amino acid is mapped uniquely to a hash value

**Table S2. aaHash Level 2 hashing scheme**

$$\Sigma_2 = \{A, (R, Q, K), N, (D, E), C, G, H, (I, L, M, V), (F, W, Y), P, (S, T)\}$$

Defines the aaHash hashing scheme in which amino acids are grouped if their pairwise BLOSUM62 substitution score is ≥ 1. Amino acids contained in the parenthesis are hashed to the same values.

**Table S3. aaHash Level 3 hashing scheme**

$$\Sigma_3 = \{(A, S, T), (R, Q, K), (N, D, E), C, G, H, (I, L, M, V), (F, W, Y), P\}$$

Defines the aaHash hashing scheme in which amino acids are grouped if their pairwise BLOSUM62 substitution score is ≥ 0. Amino acids contained in the parenthesis are hashed to the same values.

**Table S4. Benchmarking commands and parameter settings for human genome completeness assessment comparisons against state-of-the-art**

| Tool | Command |
|---|---|
| **aaKomp** | run-aakomp \ <br>--db-dir /path/to/busco_downloads \ <br>--reference /path/to/busco_downloads/primates_odb12.faa \ <br>--input assemblyN.fasta \ <br>--track-time \ <br>-o aakomp_assemblyN \ <br>--threads 48 \ <br>-H 9 \ <br>-k 9 \ <br>-l 0.7 |
| **BUSCO**(Manni et al. 2021) | /usr/bin/time -pv -o busco_assemblyN.time \ <br>busco \ <br>-i assemblyN.fasta \ <br>-m genome \ <br>-l primate_odb12 \ <br>-c 48 \ <br>-o busco_assemblyN |
| **Compleasm**(Huang and Li 2023) | /usr/bin/time -pv -o compleasm_assemblyN.time \ <br>compleasm run \ <br>-l primate_odb12 \ <br>-L /path/to/busco_downloads \ <br>-t 48 \ <br>-a assemblyN.fasta \ <br>-o compleasm_busco_assemblyN |

**Table S5. aaKomp scores across *k*-mer sizes on T2T-CHM13 human reference genome using the 'primates_odb12' miBf database.**

| *k*-mer size | aaKomp Score |
|---|---|
| 7 | 83.27 |
| 8 | 93.33 |
| 9 | 94.03 |
| 10 | 93.38 |
| 11 | 92.18 |

**Method S1. Running and benchmarking procedures for aaKomp, BUSCO, and Compleasm**
For both BUSCO and Compleasm, the 'primate_odb12' and 'actinopterygii _odb12' ortholog database was downloaded ahead of time. miBf databases for aaKomp were constructed from the corresponding reference proteomes. Both the BUSCO ortholog database and miBf database were reused across all experiments to ensure consistency. To enable fair benchmarking of runtime and performance, the time required to download BUSCO data and build miBf was excluded from runtime measurements. All tools were run using their default parameters unless otherwise specified and benchmarked using '/usr/bin/time' (S2 Table). For aaKomp, the driver script includes an internal '--track-time' option, which utilizes '/usr/bin/time' to record runtime and resource usage.

**Discussion S1. Impact of reference database complexity on computational performance and sensitivity.**
We also observed that performance varies based on the size of the chosen reference database, with the human proteome requiring more time and memory than the more compact 'primates_odb12' set. This increase reflects the inherent computational demands of querying a larger database because as the number of reference sequences grows, there are more potential *k*-mer matches to evaluate, which naturally impacts both run time and memory usage. The human proteome contains 20,663 protein entries—nearly double the 11,834 consensus sequences in 'primates_odb12'—making it a more demanding dataset to query.

**Supplementary References**

Huang, N., and Li, H. 2023. compleasm: a faster and more accurate reimplementation of BUSCO. Bioinformatics **39**(10): btad595. doi:10.1093/bioinformatics/btad595.

Manni, M., Berkeley, M.R., Seppey, M., Simão, F.A., and Zdobnov, E.M. 2021. BUSCO Update: Novel and Streamlined Workflows along with Broader and Deeper Phylogenetic Coverage for Scoring of Eukaryotic, Prokaryotic, and Viral Genomes. Molecular Biology and Evolution **38**(10): 4647–4654. doi:10.1093/molbev/msab199.
